# Subthalamic nucleus mediates the modulation on cocaine self-administration induced by ultrasonic 3 vocalizations playback in rats

**DOI:** 10.1101/450437

**Authors:** Montanari Christian, Giorla Elodie, Pelloux Yann, Baunez Christelle

**Affiliations:** Institut de Neurosciences de la Timone, UMR 7289 CNRS & Aix-Marseille Université, 27 boulevard Jean Moulin, 13005 Marseille, France.

**Keywords:** addiction, basal ganglia, conditioned place aversion, conditioned place preference, social influence, sucrose

## Abstract

Drug intake is known to be under the influence of social context. We have recently shown that presence of a peer influences drug intake in both rats and humans. Whether or not social acoustic communications between the peers play a role during cocaine or sucrose self-administration (SA) was investigated here, using playback of ultrasonic vocalizations (USV) at 50- and 22-kHz, conveying respectively positive and negative internal affective states in adult rats. To assess the neurobiological substrate of a potential USV influence on drug and food intake, we tested the effects of subthalamic nucleus (STN) lesions, given its role in emotional and motivational processes. In sham-control rats, playback of USV associated with positive affective states induced long-term decreased cocaine consumption, while USV associated with negative affective states induced short-term increase. Interestingly, no effect of USV playback was observed on sucrose intake, whatever the frequency. STN lesions abolished the influence of USV on cocaine intake, highlighting the influence of STN in emotional processes induced by USV emitted by a peer. These results show how acoustic social communication is important to regulate drug intake in rats and how STN modulation could interfere with addiction processes.

## Introduction

Drug use disorders are a major worldwide public health problem in our society (APA 2013). Recently it was demonstrated that the so-called “proximal social factors”, that are the social factors present at the time of drug exposure such as the presence and behavior of peers, play an important role in both human drug consumption and in rats self-administering drugs (Strickland & Smith 2013 for review; Giorla et al. 2018). Epidemiologic studies have shown that people associated with peers who use drugs have higher probabilities to use drugs than people not associated with drug users (Bahr, Hoffmann, & Yang 2005). Moreover in a semi naturalistic bar setting (Larsen et al. 2009) and in a real bar study (Larsen et al. 2012) it was shown that subjects in company of a heavy-drinking peer consumed more alcohol than subjects in company of a light-drinking peer or an abstinent peer. In the rat, the peer presence influences drug intake in a different manner if the peer is abstinent or is also taking drugs (see Strickland & Smith 2013 and Discussion). Abuse of food, particularly of highly palatable sweet food rich in sugars, is another serious burden of our western countries (for reviews see Ahmed, Guillem, & Vandaele 2013 and DiNicolantonio, O’Keef, & Wilson 2017). Although there is no explicit definition of “proximal social factors” in eating-related behaviors, it is well-known that presence and behavior of peers influence also our eating behavior, a daily activity often shared with others. The probability to become obese is higher if a friend, a sibling or a spouse has become obese (Christakis & Fowler 2007). Moreover, in presence of a non-eating observer, people eat less than when alone (Roth et al. 2001), and when people eat together their intake differs from when alone and depends on the others intake (Herman 2015). The social facilitation phenomenon of eating behavior is well-known in different animal species such as rats (Harlow 1932), chickens (Tolman 1965), dogs (Ross & Ross 1949) and monkeys (Harlow & Yudin 1933). Gardner & Engel (1971) showed that a rat increases its lever-pressing behavior for standard laboratory food in presence of a peer having access to food. No further studies evaluated the effect of the presence and behavior of a peer on food self-administration (SA), especially palatable sweet food that, like drugs, can induce a loss of control over consumption in both animals and humans (Ahmed, Guillem, & Vandaele 2013; DiNicolantonio, O’Keef, & Wilson 2017). Given that the influence of proximal social factors in animals has been only recently studied, a few questions remain to be addressed, such as their influence on both drug and food SA and the ways of communication used between peers to influence each other.

Adult rats use 50- and 22-kHz ultrasonic vocalizations (USV) to respectively communicate to peers their positive and negative affective internal states (Brudzynski 2013). This acoustic communication could thus play an important role in the effects mediated by the peer presence on drug and food intake. Although recording USV of rats self-administering drugs and palatable food as sucrose is common to assess the emotional status of the animals (Browning et al. 2011; Barker, Simmons, & West 2015), no study has ever evaluated the effects of USV playback on drug and food intake, albeit other recent studies have shown that USV playback in rats is an essential tool to investigate different human brain disorders, especially those involving social communication deficits (Wöhr & Schwarting 2017). Moreover there are no studies regarding the neurobiological basis of the proximal social factor’s effects on drug and food consumption. However, one can hypothesize that emotional and motivational processes may interact in the influence of proximal social factors on goal-directed behaviors. Therefore, the subthalamic nucleus (STN), a cerebral structure belonging to the basal ganglia, could represent an interesting candidate because it was previously shown to modulate affective responses to emotional stimuli (Pelloux et al. 2014) and its inactivation reduces motivation for cocaine while increasing that for food only in a Progressive Ratio schedule of reinforcement but not in standard SA conditions (Baunez et al 2005; Rouaud et al 2010). In this study we thus aimed at evaluating the effects of 50- and 22- kHz USV playback on both cocaine and sucrose SA, and in both sham-control and STN-lesioned rats.

## Material and methods

### Animals and Surgery

A total of 37 male Lister Hooded rats (Charles River Laboratories, Saint-Germain-sur-l’Arbresle, France) weighing ~290 g at their arrival were used. Animals were housed in pairs, maintained in animal facility and subjected to intra-jugular implantation of catheters and/or STN ibotenic excitotoxic lesions (rats underwent oral sucrose SA were only subjected to STN lesions) as previously described (Baunez et al. 2005; Pelloux et al. 2014; see Supporting Information). All the rats had free access to standard laboratory food pellets (Scientific Animal Food and Engineering, Augy, France) and water, except during the experimental sessions. All animal procedures were approved by local ethic committee and the University of Aix-Marseille (#3129.01), and were in accordance with the European Community Regulations for Animal Use in Research (CEE N° 86/609) and the French regulation (Decree 2010-118).

### Experiments overview

In total, 4 different experiments were performed: 1) evaluation of the effects of 50- and 22-kHz USV playback on cocaine SA; 2) evaluation of the effects of 50- and 22-kHz USV playback on sucrose SA; 3) assessment of the rewarding properties of the 50-kHz USV using the Conditioned Place Preference (CPP) paradigm; 4) assessment of the aversive properties of the 22-kHz USV using the Conditioned Place Aversion (CPA) paradigm. To limit the number of animals, the rats that first underwent the cocaine SA procedure were also tested in the CPA paradigm, and the rats tested in the CPP paradigm were subsequently used for the sucrose SA experiment (except 3 sham-control rats used only for the CPP paradigm). Between the two experimental procedures all animals were maintained for 2-3 months in standard housing conditions, without any exposure to cocaine, sucrose or USV playback. The rats were randomly allocated into the different experimental groups when the study was designed with the aim to use the smallest number of animals sufficient to consider the results reliable (*cocaine SA/CPA*: sham-control rats, N=6; STN-lesioned rats, N=9; *CPP*: sham-control rats, N=11; STN-lesioned rats, N=9; *sucrose SA*: sham-control rats, N=8; STN-lesioned rats, N=9).

### Playback display of 50- and 22-kHz USV over cocaine and sucrose SA sessions

*Apparatus*: four standard SA chambers (Camden Inst, 24- cm length, 25- cm width, and 26-cm height) placed in sound- and light attenuating cubicles were used. Each chamber was equipped with two retractable levers positioned on the right-hand wall, two white stimulus lights (one above each lever), an extractable dipper for the delivery of liquids in the food magazine (placed between the 2 levers), and a white house light located in the ceiling. Acoustic stimuli were presented through ultrasonic loudspeakers (Ultrasonic Dynamic Speaker Vifa, Avisoft Bioacoustics, Glienicke, Germany) placed in an opening in the middle of the chambers ceiling (see Supporting Information).

*Procedures*: testing began 10 days after the surgery. Different animals were used to test the USV playback on cocaine and sucrose SA. Daily 1h sessions took place during the dark phase, between 09:00 and 17:00 hours. Immediately before the start of sessions, catheters of rats that were self-administering cocaine were connected – through infusion lines and liquid swivels - to cocaine syringes positioned on motorized pumps (Razel Scientific Instruments, St. Albans, VT, USA) placed outside of the chambers. At the start of each session a white house light was turned on for the duration of the session and two lever were extended. Depending on the experiment, pressing the active lever (counterbalanced between rats) delivered either an intravenous infusion of cocaine (250µg/90µl/5s) or a 10% solution of sucrose accessible in the dipper (300µl/dipper/5s) and switched on the white cue light above the active lever during the reward delivery; pressing the inactive lever was recorded but had no consequence. A 20-s time-out period was imposed (and maintained during all the phases of the experiment), during which any further lever press was recorded as perseverative response but had no consequence. Few sessions were run under a continuous schedule of reinforcement (FR1; every lever press on the active lever was reinforced). Then progressively over days, the number of lever presses required to obtain a reward was increased to FR2 and FR5 (the 2^nd^ and the 5^th^ lever press was respectively reinforced). When rats reached a stable cocaine or sucrose SA (baseline condition: ≤20% of variability in the number of rewards for 5 consecutive days), they were exposed – for 3 different blocks of 5 consecutive days and in the following order – to the playback of 50-/22-kHz USV, background noise, and 22-/50-kHz USV, counterbalancing between rats the starting order of the USV (50- *versus* 22-kHz). During the playback sessions, the first lever press of each FR5 ratio resulted in the playback of the appropriate file (1.5 sec length) and the lever pressing counting was not interrupted during the playback (see Acoustic Stimuli section and Supporting Information for further details on the USV used and the playback procedure). The background noise was used as control stimulus of possible unspecific effects not due to the communicative function of the USV. The chambers, pumps and loudspeakers were controlled by a home-built interface and by a home-written PC-program (built and written by Y. Pelloux).

### Assessment of USV emotional properties through the Conditioned Place Preference/Aversion (CPP/CPA) paradigm

The rewarding and aversive properties of the USV were assessed using the CPP/CPA paradigm (for a review see Tzschentke 2007).

*Apparatus*: As previously described (Rouaud et al. 2010), a gray Perspex box (70 cm length × 30 cm width × 40 cm height) divided in two main lateral compartments (30 × 30 × 40 cm each) with a small intermediate chamber (10 cm in length) was used for the experiments. The two lateral compartments differed only in the spatial organization of two vertical columns (i.e. conditioned stimulus). Acoustic stimuli were played back through the same ultrasonic loudspeakers used for the SA sessions, placed above the middle of the lateral compartments of the apparatus. A camera (PNP-90/TL, Swann Communications Ltd, Central Milton Keynes, UK) connected to a Computer Video Interface (EzCAP USB Video Grabber, EZCAP.TV, Newton Stewart, UK) was mounted above the apparatus for video recording.

*Procedures:* different animals were used for the CPP and the CPA experiment. Rats that underwent the CPP experiment were exposed to the 50-kHz USV, while the ones that underwent the CPA experiment were exposed to the 22-kHz USV. Briefly, on the preconditioning day, rats explored the whole apparatus for 15 min and the time (in sec) spent in each compartment was measured via the video recording. From the following day and for 8 consecutive days (conditioning phase: 30 min/session/day), animals were confined - on alternate days- one day in one of the two lateral compartments and exposed to the USV (50- or 22- kHz USV), and the day after in the other lateral compartment and exposed to the background noise [counterbalancing between rats the lateral compartment paired with the USV and the starting order of the acoustic stimuli (USV *vs* background noise)]. The same acoustic files (1.5 sec length) played back during the SA sessions were displayed during each conditioning session, every 5 minutes for a total of 5 playbacks/session (see Acoustic Stimuli section and Supporting Information for further details on the USV used and the playback procedure). The day after the end of the conditioning, on the testing day, rats were again let free to explore the apparatus for 15 min without any playback and the time spent in each compartment was measured via the video recording, as in the pre-conditioning day. The rewarding or aversive properties of the USV were inferred by the measurement of the time spent during the test in the compartment previously paired with the USV in comparison with the time spent in the same compartment in the preconditioning day.

### Acoustic Stimuli

Animals were tested for their auditory capacity before the beginning of the experiments, after recovery from surgery; all rats showed an excellent startle response after a sudden sound stimulation and then no rat was excluded from the experiments due to hearing deficits. The USV displayed during the experiments were recorded from 2 adult male Lister Hooded rats, unfamiliar to the tested rats and not included in any experimental procedure. A long (1.5s) 22-kHz USV call was recorded from a rat that received an electric foot shock (0.5 mA for 2s) in a SA chamber before the recording. For playback a 50-kHz USV, the length had to be identical to the 22-kHz USV. Since such long 50-kHz USV do not exist, a series of 50-kHz calls (total file length: 1.5s; total calling time: 0.45s) was recorded from a rat during the exploration of a SA chamber containing food, water and scents from its cage mate after depriving him overnight from all of the above reported stimuli (see Supporting Information for further details on the USV used).

A Condenser Ultrasound Microphone (CM16/CMPA-48, Avisoft Bioacoustics) and its Computer Audio Interface (QUAD-CAPTURE USB 2.0 Audio Capture, Roland Corporation, Los Angeles, CA, U.S) were used to record the USV with a sampling rate of 192 kHz in 16-bit format using Avisoft RECORDER (Version 4.2, Avisoft Bioacustics).

### Histology

See Supporting Information

## Statistical analysis

The statistical analyses have been performed using SPSS 20 software and all variables were expressed as mean number ± SEM.

*Playback display of 50- and 22-kHz USV over cocaine and sucrose SA sessions*: in both sham-control and STN-lesioned rats the number of rewards reached during the sessions was analyzed via a two-way repeated measures ANOVA with playback condition (4 levels: baseline, background noise, 50- and 22- kHz USV) and session (5 levels: five days for each playback condition) as within-subject factors. The average of the baseline condition was also compared with the number of rewards obtained during the 1^st^ day of the other playback conditions, through a one-way repeated measure ANOVA. Significant ANOVA effects (p ≤0.05) were followed by post hoc Fisher’s LSD tests. No statistical methods were used to predetermine sample size, but the samples are comparable to those reported in previous studies. However, power analysis (1-group, 2-tails and equal variance model) was performed, using the Free Online Power and Sample Size Calculator available at powerandsamplesize.com, to confirm that our sample sizes were sufficient to detect reliable changes in cocaine intake under USV playback influence. Comparisons between groups on the number of cocaine/sucrose rewards reached by rats during the 1^st^ day of playback and for the average of each 5-day block, were made using independent t-test (2-tails).

*Assessment of the USV emotional properties through the Conditioned Place Preference/Aversion (CPP/CPA) paradigm:* we calculated the preference/avoidance score as the difference between the time (in sec) spent on the testing day in the compartment previously associated with the USV and the time spent in the same compartment on the preconditioning day. A positive score indicates a preference, while a negative score an avoidance, for the compartment previously paired with the USV. The conditioning effect was evaluated within each group via the non-parametric Wilcoxon signed-rank test for paired samples, while differences between groups were assessed through the non-parametric Mann-Whitney U test for independent samples.

## Results

### Histology

Five rats (2 and 3 from the cocaine SA/CPA and sucrose SA/CPP experiments respectively) were excluded from the STN-lesioned groups results and statistical analyses due to lesions either unilateral, too restricted or outside the STN. The sites and extent of lesions were characterized by a neuronal loss and associated gliosis (see Fig. 1 for a representative intact and lesioned STN).

**Figure 1.**
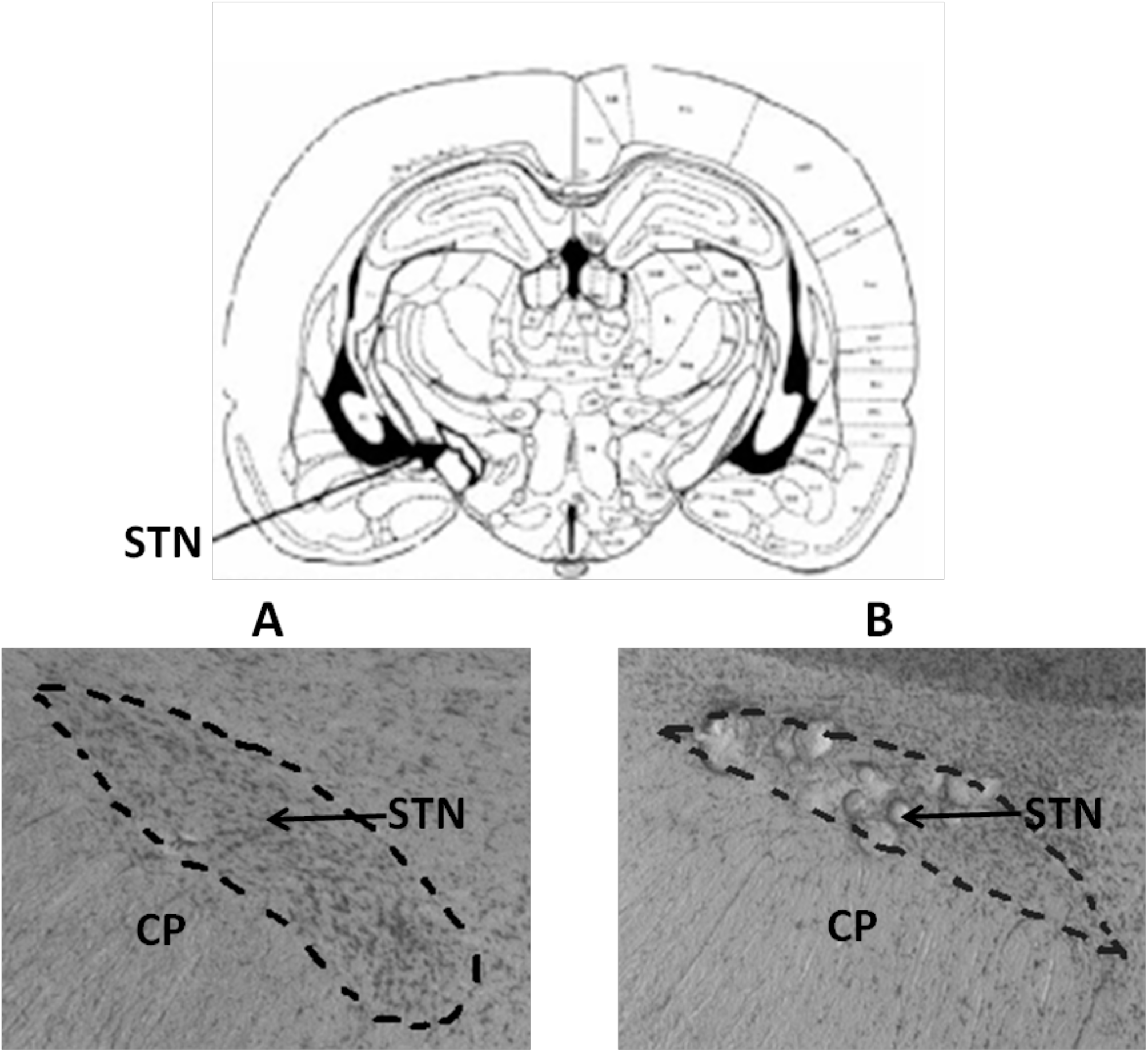
Frontal STN sections stained with cresyl violet. *Top panel:* coronal section of the rat brain at the level of the STN (left STN indicated by the arrow), from Paxinos & Watson 2005. *Bottom panels:* the STN is delineated by the dashed lines in a representative sham-operated rat (A) and a representative STN-lesioned rat (B). The lesions were characterized by neuronal loss, gliosis and calcifications. CP: cerebral peduncle; STN: subthalamic nucleus.

### Effects of 50- and 22-kHz USV playback on cocaine SA sessions

Thirteen rats were included in the analysis (sham-control group: N=6; STN-lesioned group: n=7).

*Sham-control rats*: the analysis of the effects of the USV playback across all sessions revealed a significant effect of playback condition [ANOVA: F(3,15)=6.711, p=0.020], but not of session [ANOVA: F(4, 20)=3.365, p=0.060] nor playback condition × session interaction [ANOVA:F(12,60)=2.269; p=0.118]. Post hoc comparisons between the different playback conditions, illustrated in Fig. 2*a* and in Fig. 2*b* as averaged by condition, indicated that the playback of background noise did not affect cocaine intake compared with baseline (p=0.240), whereas the playback of the 50-kHz USV induced a significant decrease in the number of cocaine injections compared to baseline (p=0.048, power=0.99), background noise (p=0.034, power=0.99) and 22-kHz USV (p=0.028, power=1), and the 22-kHz USV had no significant effect when compared to baseline and background noise (p=0.244; p=0.094 respectively). Interestingly, when the analysis only included the 1^st^ day of each playback condition, as illustrated in Fig 2*c*, there was a significant playback effect [ANOVA: F(3,15)=9.802, p=0.001] and post-hoc comparisons revealed that baseline and background noise did not affect cocaine intake differently (p=0.723), and that both 50- and 22- kHz USV had a significant effect, the first decreasing cocaine intake (p=0.046 with power=0.60 and p=0.041 with power=0.71, from baseline and background respectively), while the second increased it (p=0.034 with power=0.92 and p=0.024 with power=0.94, from baseline and background respectively), leading to a significant difference between 50- and 22 kHz USV (p=0.019, power=1).

**Figure 2.**
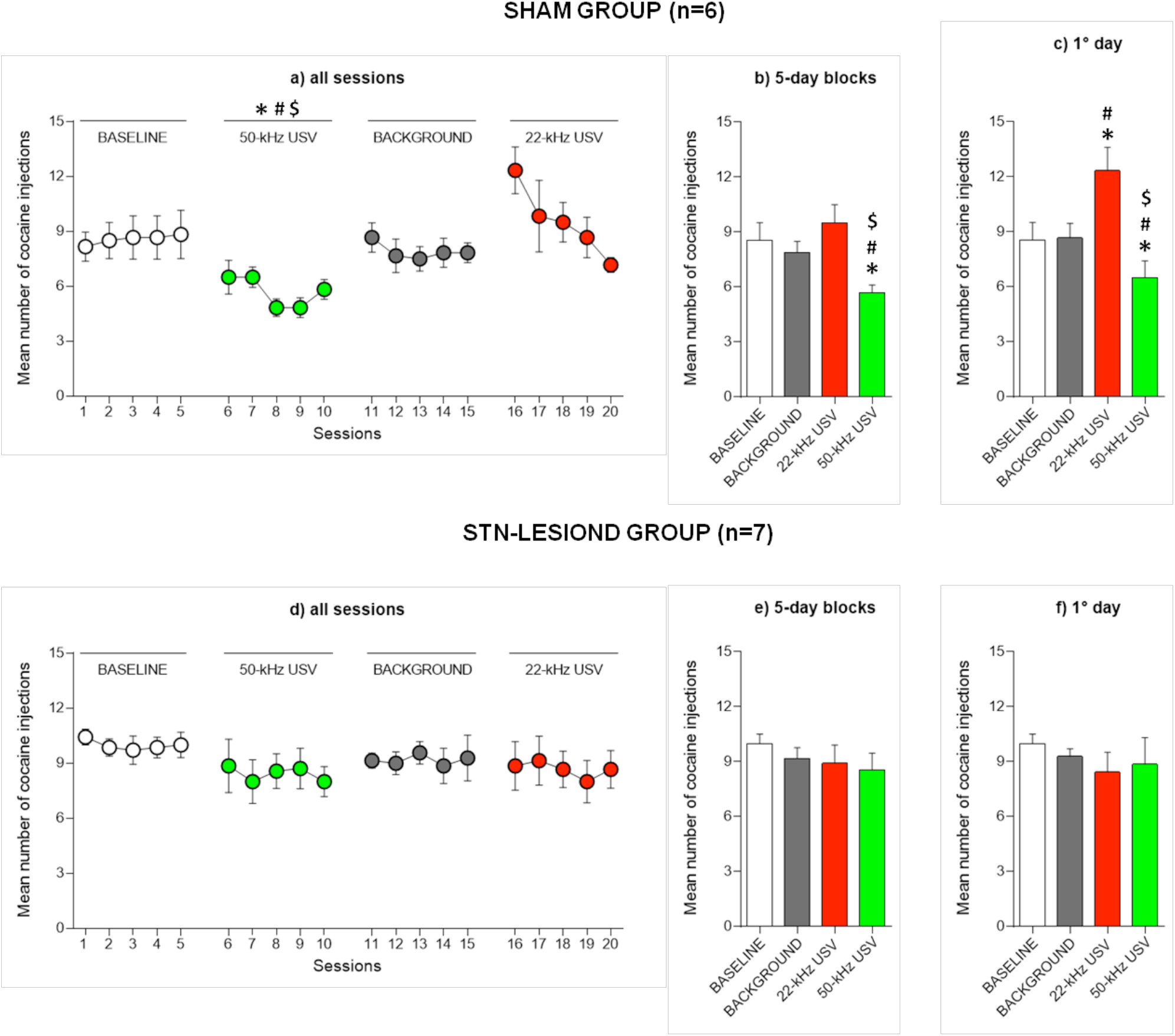
Effects of 50-kHz USV, 22-kHz USV and background noise playback on the mean number of cocaine injections obtained. The graphs represent the mean number (± SEM) of cocaine injections (250µg/90µl/5s) taken from sham-control (N=6; *a-c*) and STN-lesioned rats (N=7; *d-f*) under the various conditions of playback during all the sessions (*a,d*), during each 5-day block (*b,e*) and during the 1^st^ day of playback (*c,f*). *: p≤ 0.05 compared with baseline; #: p≤ 0.05 compared with background noise; $: p ≤ 0.05 compared with 22-kHz USV.

*STN-lesioned rats*: in this group the number of cocaine injections was not affected by the different playback conditions (Fig. 2*d-f*). As shown in Fig. 2*d* and in Fig. 2*e* as averaged by condition, the analysis across the sessions did not indicate a significant effect of playback condition [ANOVA: F(3,18)=0.800, p=0.468], session [ANOVA: F(4,24)=0.371, p=0.772] or playback condition × session interaction [ANOVA: F(12,72)=0.600, p=0.624]. As shown in Fig. 2*f*, no playback condition effect was further found on the 1^st^ day of playback [ANOVA: F(3,18)=0.653, p=0.513]. Comparisons between groups showed that sham-control rats, in comparison to STN-lesioned rats, took a higher number of cocaine injections during the 1^st^ day of the 22-kHz USV playback [t(11)=2.322; p=0.42], and a lower number of injections during the 50-kHz 5-day block [t(11)=-2.837; p=0.021].

### Effects of 50- and 22-kHz USV playback on sucrose SA sessions

Fourteen rats were included in the analysis (sham-control group: N=8; STN-lesioned group: N=6).

*Sham-control rats*: the number of sucrose rewards obtained was stable regardless of playback condition and session (Fig. 3*a-c*). The analysis of the number of sucrose rewards earned across all sessions did not indicate any significant effect of playback condition [ANOVA: F(3,21)=0.898, p=0.433], session [ANOVA: F(4,28)=1.315, p=0.289] or playback condition × session interaction [F(12,84)=0.898, p=0.530], as shown in Fig. 3*a* and in Fig. 3*b* as averaged by condition. As shown in Fig. 3*c*, no significant playback condition effect was further found in the analysis of the 1^st^ day of playback [ANOVA: F(3,21)=1.376, p=0.284].

**Figure 3.**
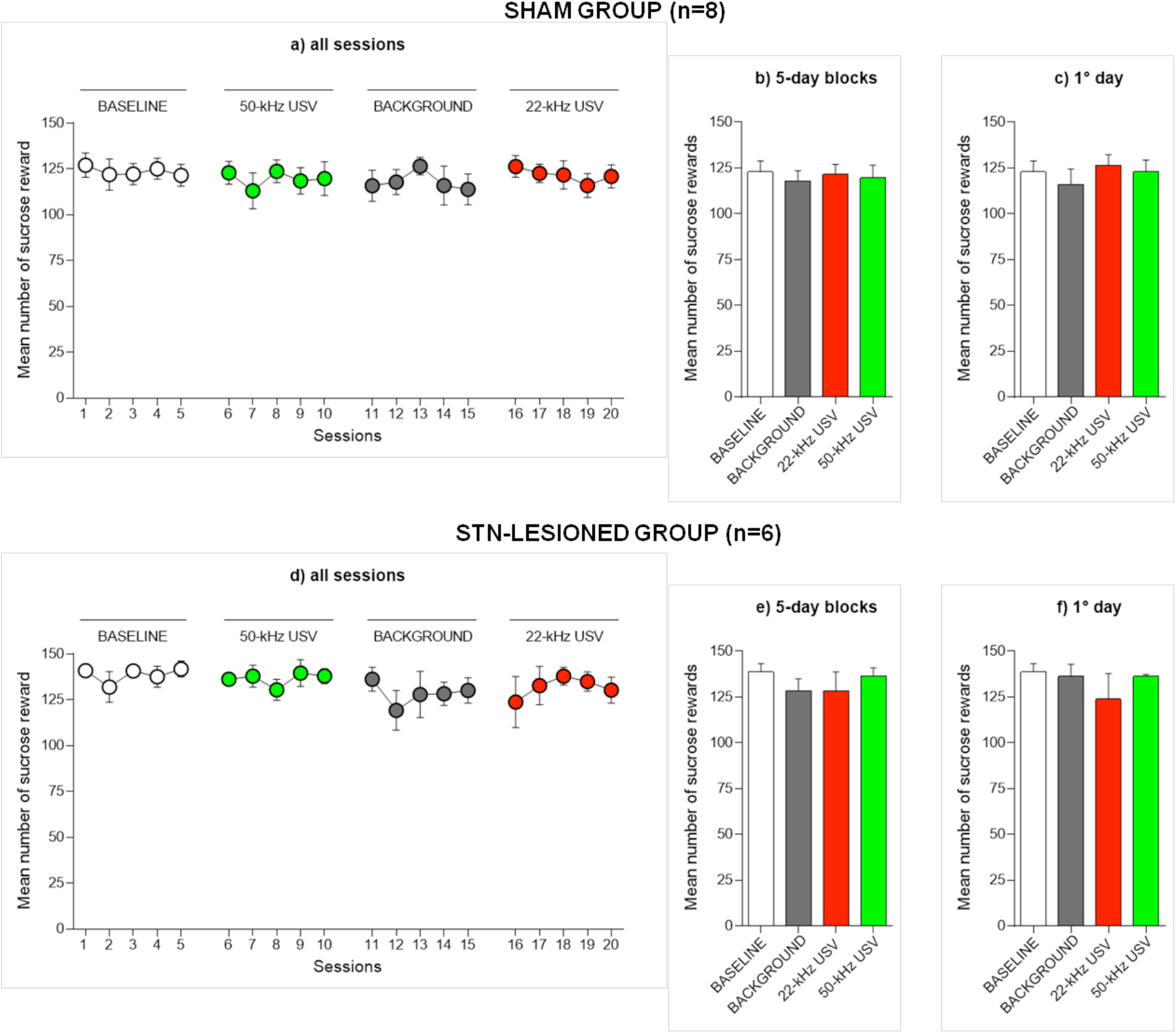
Effect of 50-kHz USV, 22-kHz USV and background noise playback on the mean number of sucrose rewards obtained. The graphs represent the mean number (± SEM) of dipper activations giving access to a 10% liquid solution of sucrose (300µl/dipper/5s) taken from sham-control (N=8; *a-c*) and STN-lesioned rats (N=6; *d-f*) under the various conditions of playback during all the sessions (*a,d*), during each 5-day block (*b,e*) and during the 1^st^ day of playback (*c,f*).

*STN-lesioned* rats: the number of sucrose rewards in STN-lesioned rats was not modulated by the different acoustic stimuli played back (Fig. 3*d-f*). As shown in Fig. 3*d* and in Fig. 3*e* as averaged by condition, no significant effect of playback condition [ANOVA: F(3,15)=1.802, p=0.190], session [ANOVA: F(4,20)=1.065, p=0.383] or playback condition × session interaction [ANOVA: F(12,60)=0.913, p=0.458] was found. As illustrated in Fig. 3*f*, the absence of a significant playback condition effect was confirmed in the analysis of the first 1^st^ day of playback [ANOVA: F(3,15)=1.246, p=0.328].

### Assessment of the rewarding and aversive properties of the 50- and 22-kHz USV

For the CPP and CPA experiments, 17 sham-control (N=11 and 6 respectively) and 13 STN-lesioned rats (N=6 and 7 respectively) were included in the analysis. As shown in Fig. 4, only in the sham-control group, playback of the 50- and 22-kHz USV respectively induced a preference and an avoidance towards the environment previously paired with the USV. As shown in Fig. 4*a*, the playback of the 50-kHz USV induced a conditioning effect for the compartment paired with the USV in sham-control but not in STN-lesioned rats (Wilcoxon signed-rank test: p=0.026 and p=0.400, respectively), leading to a significant difference between the two groups (Mann-Whitney U test: p=0.027). Similarly, and as shown in Fig. 4*b*, the playback of the 22-kHz USV induced a conditioning effect for the compartment paired with the call in sham-control but not in STN-lesioned rats (Wilcoxon signed-rank test: p=0.028 and p=1.000, respectively), leading again to a significant difference between the two groups (Mann-Whitney U test: p=0.022).

**Figure 4.**
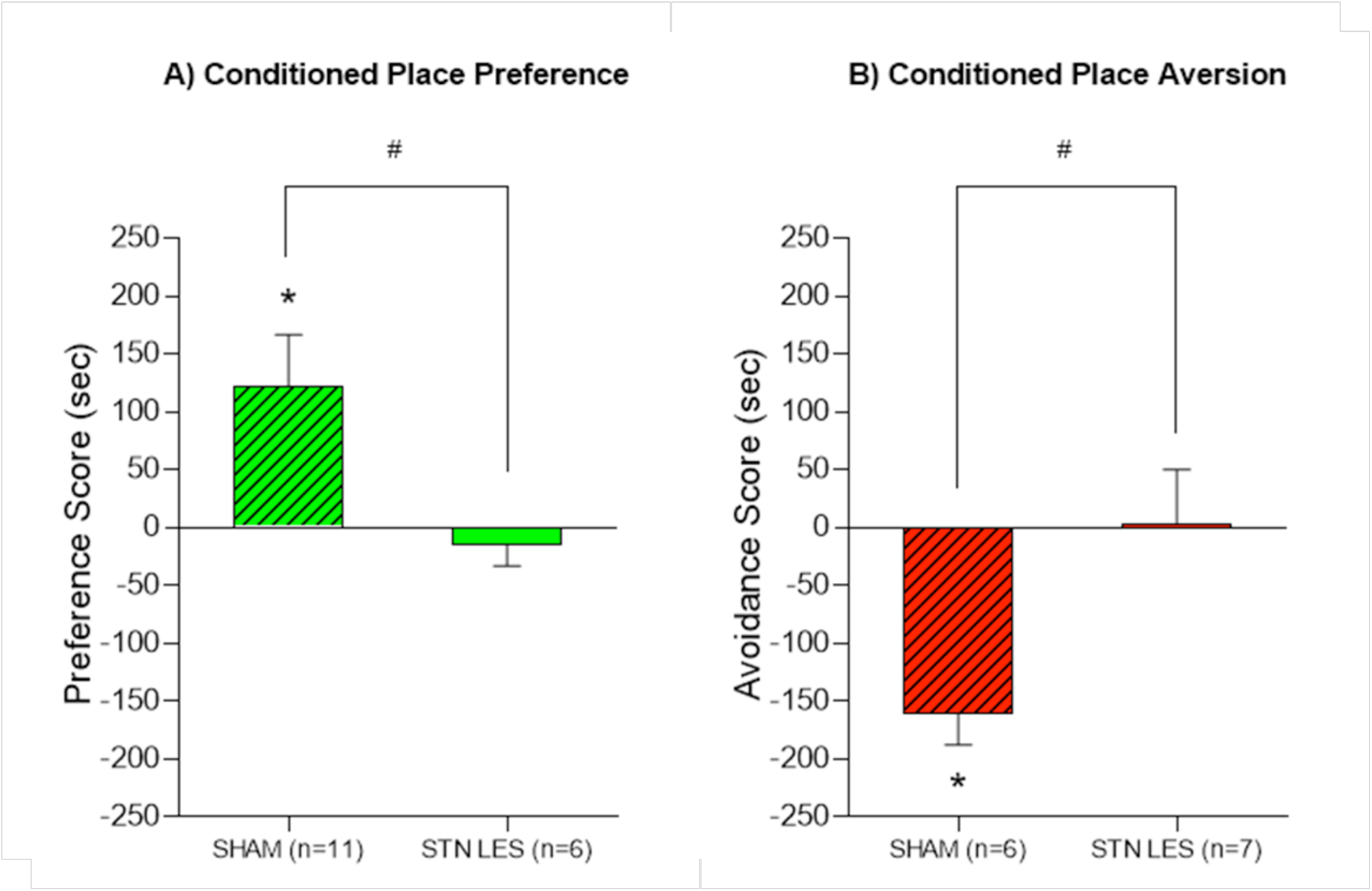
Conditioned Place Preference and Conditioned Place Aversion for the compartment respectively paired with the 50- and 22-kHz USV. The figure shows the mean (±SEM) preference and avoidance score, respectively for the 50- and 22-kHz USV, calculated as the time (sec) spent during the test day in the compartment previously associated with the USV minus the time spent in the same compartment on the preconditioning day. *a:* preference score for the compartment paired with the 50-kHz USV in both sham-control(N=11) and STN-lesioned rats (N=6) [conditioning effect: *, p≤ 0.05; group effect: #, p≤ 0.05]. *b:* avoidance score for the compartment paired with the 22-kHz USV in both sham-control (N=6) and STN-lesioned rats (N=7) [conditioning effect: *, p≤ 0.05; group effect: #, p ≤0.05].

## Discussion

### Affective meaning of USV and consequences on cocaine intake

We have shown here that in sham-control rats the playback of 50-kHz USV induced a decreased intake of cocaine, while the playback of 22-kHz USV increased it transiently. The 50-kHz calls may have acted as an alternative reward able to reduce cocaine intake, as it was previously reported for other rewards. It was shown indeed that four regular food pellets placed in the self-administration chambers reduced heroin intake in rats fed *ad libitum* (Lenoir & Ahmed 2008). Moreover in a mutually exclusive choice procedure, majority of rats fed *ad libitum* choose to self-administer sugar solutions over cocaine and other drugs (see Ahmed, Guillem, & Vandaele 2013 for a review). The presence of a peer has also been shown to be rewarding for rats, given that they develop a social interaction-induced CPP (Calcagnetti & Schechter 1992; Giorla et al. 2018) and learn to press a bar for a physical contact with another rat or only its view (Angermeier 1960). Interestingly, in the SA studies that evaluated the effects of peers on drug intake, rats tested with a non-self-administrating peer took less cocaine than rats tested with a peer also taking cocaine or rats tested alone (Smith 2012; Robinson et al. 2016; Giorla et al. 2018). Here we have shown that the playback of 50-kHz calls induced a CPP, confirming the positive meaning of the 50-kHz USV (Brudzynski 2013). This is particularly in line with the highest positive affective states conveyed by the frequency-modulated (FM) and trills calls (Burgdord et al. 2008; Burgdorf & Moskal 2010), of which our file is in majority composed (see Acoustic Stimuli section and Supporting Information for further details on the USV used). Our CPP result is also in agreement with the fact that rats can self-administer playback of 50-kHz USV (Burgdorf et al. 2008), and that 50-kHz USV playback induces dopamine release (Willuhn et al. 2014) and neuronal activation (Sadananda, Wohr, & Schwarting 2008) in the nucleus accumbens, these latter results being interpreted as a stimulation of the reward system. If we then consider the apparently rewarding properties of the 50-kHz USV and the ability of other rewards to reduce cocaine intake, it is not surprising that also the 50-kHz calls were able to do so. It was also well documented that 50-kHz USV serve an important affiliative pro-social function (Wöhr et al. 2016; Wöhr & Schwarting 2017). Consequently it could be possible that rats decreased their cocaine intake because they were approaching the loudspeaker and looking for a peer. However, given that in a previous report rats showed an approach behavior towards the source of the 50-kHz USV playback only during the first USV exposure (Wöhr & Schwarting 2012), we should not expect it to last as long as 5 consecutive SA sessions. Moreover, our data obtained with the CPP (a classical and well established paradigm to study the rewarding properties of drug and non-drug stimuli; see Tzschentke 2007 for a review), showed that rats, during the test day, kept approaching the compartment associated with the USV in absence of their playback. Considering the 50-kHz USV are uttered during – mostly positive – various social interactions (Burgdorf et al. 2008), but also after exposure to food and drugs (Browning et al. 2011; Barker, Simmons, & West 2015), or in mildly aversive situations as a short social isolation (Wöhr et al. 2007), we can reasonably suppose that the social and rewarding aspects of the 50-kHz USV could overlap in some circumstances, while in other situations one aspect could prevail on the other. Further research is necessary to investigate this point.

In another hand, the playback of the 22-kHz USV probably increased cocaine intake through the induction of an aversive state, in line with a series of results indicating higher levels of cocaine intake after exposure to stressful stimuli in rats (Goeders & Guerin 1994). In support of this hypothesis, the long 22-kHz call used here induced a CPA, confirming the aversive meaning of the 22-kHz USV, in particular of the long ones (Brudzynski et al. 1993; Brudzynski 2013; see Acoustic Stimuli section and Supporting Information for further details on the USV used). Moreover previous findings indicated that rats avoid SA of 22-kHz USV playback (Burgdorf et al. 2008), and that their playback induces neuronal activation in areas correlated to fear regulation such as amygdala and periaqueductal grey area (Sadananda, Wohr, & Schwarting 2008). Interestingly, starting from the 2nd day of playback, the 22-kHz USV had no effect on cocaine intake, suggesting that rats could have habituated to the aversive properties of the calls (Rankin et al. 2009). However none was observed with the 50-kHz USV, and repeated 22-kHz USV playback was able to induce – in the same animals - a CPA after 2-3 months from the end of the cocaine SA experiment, ruling out the hypothesis of an habituation process. Alternatively, the 22-kHz USV could be less negative than 50-kHz USV were rewarding, albeit in both cases we used highly affective stimuli (see above and Supporting Information). Taken all together, our results seem to suggest that the aversive properties of the 22-kHz USV were apparently maintained along the SA sessions, without however interfere with cocaine consumption, possibly because animals have quickly understood the absence of any physical threat.

### The USV-induced modulation is specific to cocaine

Our results also indicate that the USV ability to affect cocaine intake in sham-control rats is specific to the drug-related behavior, as it was not observed when animals had access to sucrose, suggesting that the rewarding properties of sucrose overcame the emotive properties of the USV. Interestingly, although the rats were fed *ad libitum*, the number of sucrose rewards was much higher than the number of cocaine infusions (122.91±5.95 sucrose dippers *versus* 8.57±0.95 cocaine infusions). Even if it can be argued that the rewarding value of a sucrose reward could be lower than a cocaine infusion, there is a large literature showing that sugar can be addictive, inducing binge-like behavior and withdrawal symptoms in rodents (DiNicolantonio, O’Keef, & Wilson 2017), and that addictive properties of sugar can be indeed stronger that those of cocaine and other drugs such as nicotine and heroin (Lenoir et al. 2007; Vandaele et al. 2016; Huynh et al. 2017). This has been recently explained by the “*delay drug reward hypothesis*” (Ahmed 2017), suggesting that the delay between an intravenous drug injection and its actions on the brain reward systems diminishes “apparently” its rewarding efficacy, as opposed to the immediate rewarding effect of sugar. In another hand it could be argued that the sucrose experiment was conducted 3 months after the rats were exposed to 50-kHz USV in the CPP experiment, exposure that may prevent the playback of USV to affect sucrose SA. However, if a former exposure to USV was sufficient to prevent them to modulate behavior, the exposure to 22- kHz USV in the cocaine SA experiment should have prevented the development of a CPA induced by 22- kHz USV playback tested 3 months later. Since CPA was observed, we can rule this hypothesis out.

### The neurobiological substrate for such modulation requires intact STN

The last noteworthy finding of our research is the inability of the 50- and 22-kHz USV playback to affect both cocaine and sucrose SA in STN-lesioned rats. It could be due to the fact that STN-lesioned rats were not able to decode the emotional properties of the USV. In fact STN-lesioned rats did not develop a preference or an aversion for the CPP/CPA compartment respectively paired with the 50- and 22-kHz USV, in agreement with their blunted emotional responses to both positive and negative stimuli previously reported (Pelloux et al. 2014). The STN has long been considered a motor structure and is the target of the surgical treatment for patients with Parkinson’s disease using the High Frequency Stimulation (HFS), also called Deep Brain Stimulation (DBS) (Limousin et al. 1995). Although the action mechanisms of the DBS are still not fully understood, they seem to partly mimic the effects of lesions of the targeted area (Gubellini et al. 2009). Interestingly STN DBS of Parkinsonian patients alters the processing of emotive information regardless the positive or negative stimulus valence, and regardless the visual or auditory sensory modality (for a review see Péron et al. 2013). Regarding the acoustic sensory modality, Parkinson’s disease patients under STN DBS show deficits in identifying different vocal emotions (Péron et al. 2010) and in contrast, decoding of emotional vocal expressions in healthy subjects involves an activation of the STN in functional magnetic resonance imaging (fMRI) studies (Péron et al. 2016). Taken all together, these human discoveries and our findings in STN-lesioned rats suggest that STN is a crucial node through which the emotional context modulates motivational processes, but also highlight possible side-effects in PD patients that should be monitored carefully. These results are also relevant for other pathologies for which STN can be a surgical target, such as Obsessive Compulsive Disorders (OCD; Mallet et al. 2008) or cocaine addiction (Pelloux & Baunez 2013). Indeed, highlighting the possible risk to impair emotions in patients treated with STN DBS should lead to a better monitoring of these patients. In another hand, this decrease in emotional functioning might explain the beneficial effects of STN DBS in OCD patients by possibly reducing a pathological over-emotionality.

### Conclusions

In conclusion we found that, in sham-control rats, 50- and 22-khZ USV are respectively rewarding and aversive. Playback of the rewarding USV leads to decreased cocaine intake, while playback of the aversive one increased it transiently. Interestingly these effects are specific to the drug-related behaviors as they were not observed when animals had access to sucrose. The neurobiological substrate for these behavioral observations seems to involve the STN since STN-lesioned rats were not able to decode the USV emotional content, as expressed by the lack of any effect in both the CPP/CPA and the cocaine SA experiments. These results demonstrate how social acoustic communication and positive contexts created either by USV or presence of a peer are important in rats to diminish drug intake and how STN modulation could help preventing detrimental effect of negative emotional and motivational contexts. These results can translate to the influence of proximal social factors on drug intake described in human drug users (Giorla et al. 2018) and could thus help improving risk reduction strategies. They are also critical for all patients that could be subjected to STN lesions or DBS for PD, OCD or even addiction, who should be monitored carefully regarding their emotional state.

Supporting Information is available online at Addict Biol’s website.

## Acknowledgements

This research was funded by CNRS, Aix-Marseille Université (AMU), the “Agence Nationale pour la Recherche” (ANR_2010-NEUR-005-01 in the framework of the ERA-Net NEURON to CB and supporting YP), the Fondation pour la Recherche Médicale (FRM DPA20140629789 to CB), and the support of the A*MIDEX project (ANR-11-IDEX-0001-02) funded by the « Investissements d’Avenir » French Government program, managed by the French National Research Agency (ANR). All the authors declare no competing interests.

## Author contribution

CM designed and performed the research, analyzed data and wrote the paper; EG performed experiments; YP designed the research and contributed to analytic tools; CB designed and supervised the research, performed surgery and wrote the paper.

